# Detection of Penicillin G Produced by *Penicillium chrysogenum* KF 425 in Vivo with Raman Microspectroscopy and Multivariate Curve Resolution-Alternating Least Squares Methods

**DOI:** 10.1101/2020.03.10.984930

**Authors:** Shumpei Horii, Masahiro Ando, Ashok Z. Samuel, Akira Take, Takuji Nakashima, Atsuko Matsumoto, Yōko Takahashi, Haruko Takeyama

**Affiliations:** Department of Advanced Science Engineering, Waseda University, 3-4-1 Okubo, Shinjuku-ku, Tokyo 169-8555, Japan; Consolidated Research Institute for Advanced Science and Medical Care, Waseda University, 3-4-1 Okubo, Shinjuku-ku, Tokyo 169-8555, Japan; Computational Bio Big-Data Open Innovation Laboratory, AIST-Waseda University, 3-4-1 Okubo, Shinjuku-ku, Tokyo 169-8555, Japan; Research Organization for Nano & Life Innovation,Waseda University, 513 Wasedatsurumaki-cho, Shinjuku-ku, Tokyo 162-0041, Japan; PRESTO, Japan Science and Technology Agency, 4-1-8 Honcho, Kawaguchi, Saitama 332-0012, Japan; Kitasato Institute for Life Sciences, Kitasato University, 5-9-1 Shirokane, Minato-ku, Tokyo 108-8641, Japan; Department of Life Science and Medical Bioscience, Waseda University, 2-2 Wakamatsu-cho, Shinjuku-ku, Tokyo 162-8480, Japan

## Abstract

Raman microspectroscopy is a minimally invasive technique that can identify molecular structure without labeling. In this study, we demonstrate in vivo detection of the bioactive compound penicillin G inside *Penicillium chrysogenum* KF425 fungus cells. Highly overlapped spectroscopic signatures acquired using Raman microspectroscopic imaging are analyzed using a multivariate curve resolution-alternating least squares (MCR-ALS) method to extract the pure spectra of individual molecular constituents. In addition to detecting multiple constituents such as proteins and lipids, we observe the subcellular localization of penicillin G like granule particle inside the fungus body. To date, there have been no reports of direct visualization of intracellular localization of penicillin G. The methodology we present in this article is expected to be applied as a screening tool for the production of bioactive compounds by microorganisms.

---

Antibiotics are indispensable for modern medicine in the treatment of diseases caused by pathogenic bacteria.^1^ In addition, these drugs have been widely used as antifungal, antiviral, and anticancer agents in humans.^2,3^ They are often discovered from secondary metabolites produced by microorganisms which are treasure box for antibiotics because they survive adaptively to a wide variety of environments.^4^ To date, 22,500 antibiotics with various pharmacological effects have been identified from microorganisms.^5^ A representative antibiotic is penicillin, which inhibits bacterial cell-wall synthesis. Penicillin was discovered from the culture broth of *Penicillium chrysogenum* by Alexander Fleming in 1928. Ernst Chain and co-workers begun to purify and study the chemistry of penicillin in 1939. Because of the poor penicillin productivity of the mold strain, 500 L of the mold filtrate per a week were necessary for a program of animal experiments and clinical trials of penicillin. This *Penicillium* strain, however, produced only traces of penicillin when grown in submerged culture. Therefore, various *Penicillium* strains were screened in a global search, and better penicillin-producing strains were obtained. Furthermore, it was found that upon exposing the culture of *P. chrysogenum* to X-rays and ultraviolet radiation, its productivity was increased still further.^6^ This strain was obtained thanks to a great amount of effort: the isolation of many wild *Penicillium* strains was required, as well as the cell culture to produce penicillin and the extraction of the penicillin, and bioassays to verify the effectiveness of the extracted compound had to be carried out. In 1942, over 10 years after its discovery, penicillin was ready for practical use. If it had been possible to detect a high-producing strain instantly, from its mold colony, without extraction and bioassay, penicillin may have been developed earlier, saving more human lives.

Raman spectroscopy can be applied to the in vivo molecular analysis of biological samples. It provides characteristic information on the molecular structure of metabolites and does not require any sample pretreatment such as dye labeling or genetic manipulation. In addition, it allows rapid and minimally invasive observation, and can be used for detection of the proteins and lipids constituting living cells.^7,8^ Moreover, Raman microspectroscopy, the combination of Raman spectroscopy with optical microscopy, allows the creation of high-spatial-resolution (∼300 nm), molecular distribution images at the single-cell level.^9^ Therefore, our research group has been investigating the application of Raman spectroscopy as a primary screening method in the search for new antibiotics, for the rapid and facile execution of investigative steps, up to and including the identification of new antibiotics. We reported the successful Raman imaging of *Streptomyces* species using confocal Raman microspectroscopy.^10^ The Raman spectroscopic images of these bacteria reveal clear and distinct spatial distributions of proteins, lipids, cytochromes, and so on. At the same time, we successfully visualized the distribution of amphotericin B (Amph-B) as a secondary metabolite. Thus, we were able to confirm that Amph-B is locally accumulated in mycelia.

However, when Raman imaging technology is applied practically to various types of microorganisms, a difficulty is encountered. Raman spectra obtained from biological systems are typically superpositions of spectra of multiple molecular species. The Raman spectra are derived from multiple constituent biomolecules, and background spectra originating from substrates and autofluorescence also contribute. If Raman bands of the specific molecule of interest do not overlap with spectra originating from other constituent molecules, it is straightforward to identify the existence of the target molecule, as already demonstrated.^10^ In contrast, if Raman bands of the target molecule overlap with those of other cellular components, which is more commonly the case, the analysis becomes much more difficult. Specifically, the overlap hinders the accurate estimation of Raman scattering intensities that originate purely from the target molecule. Therefore, a suitable analytical method is required for the separation of the spectral components of interest from those of the other cellular constituents.

Hence, to overcome this problem, we have applied a multivariate curve resolution-alternating least squares (MCR-ALS) method to decompose complicated Raman spectra into spectral components that are assignable to specific molecules.^11^ In the MCR method, the measured data is approximated as a linear combination of several spectral components. The decomposition of the spectral data is carried out via an ALS calculation. As this calculation is carried out under the constraint of all the spectral profiles and intensities being non-negative, physically and chemically interpretable solutions are preferably obtained.^11^ It should be emphasized that, rather than focusing on a few Raman bands, MCR-ALS analysis can extract components as patterns spanning the entire spectrum, so that even if some bands overlap with those of other molecular components, the spectral contribution of a particular molecule can be accurately determined. To date, MCR-ALS-based methods have been applied to the analysis of various biomaterials, as well as for the successful analysis of ingredients in pharmaceutical tablets, for the detection of marker molecules for cancer diagnosis, and for the imaging of multiple molecular components during cell division.^12–15^

In this article, we show that MCR-ALS analysis of Raman microspectroscopy results allows the detection and visualization of the production of spectroscopically overlapping secondary metabolites. A study on *Penicillium chrysogenum* is reported, and Raman mapping measurements of the fungal cells are presented. *P. chrysogenum* is known to produce penicillin G, but the Raman spectrum of penicillin G overlaps with those of proteins, which makes its analysis difficult. By using the MCR-ALS method, spectral separation of penicillin G was achieved, and localized accumulations within the cells were visualized.

## RESULTS AND DISCUSSION

### Confirmation of Penicillin G Production by *Penicillium chrysogenum* KF 425

The antibacterial activity of the mycelium extract and supernatant of *P. chrysogenum* KF 425 were tested against *K. rhizophira* NBRC 103217 and observed to have the same inhibition zone (38 mm). *P. chrysogenum* is known to produce some β-lactam antibiotics such as penicillin G, penicillin V, and cephalosporin C.^16^ Therefore, the antibiotic showed antibacterial activity against *K. rhizophira* NBRC 103217, as identified by high-performance liquid chromatography (HPLC) method. As shown in Figure 1, the penicillin G reference standard was eluted at 12.4 min. For both the cell extract samples, a penicillin G peak can be seen at the same elution time. The concentration of penicillin G produced by strain KF 425 was calculated from the standard curve based on the height of the peak in penicillin G. The penicillin G peak intensities in the mycelium extract and supernatant were 1.14 and 1.20, respectively. The corresponding concentrations of penicillin G are 22 µg/mL, in the supernatant, and 21 µg/mL, in the mycelium extract. In contrast, the non-productivity in tryptic soy broth (TSB) for penicillin G of *P. chrysogenum* KF 425 was confirmed by antimicrobial activity testing and HPLC (data not shown).

**Figure 1.**
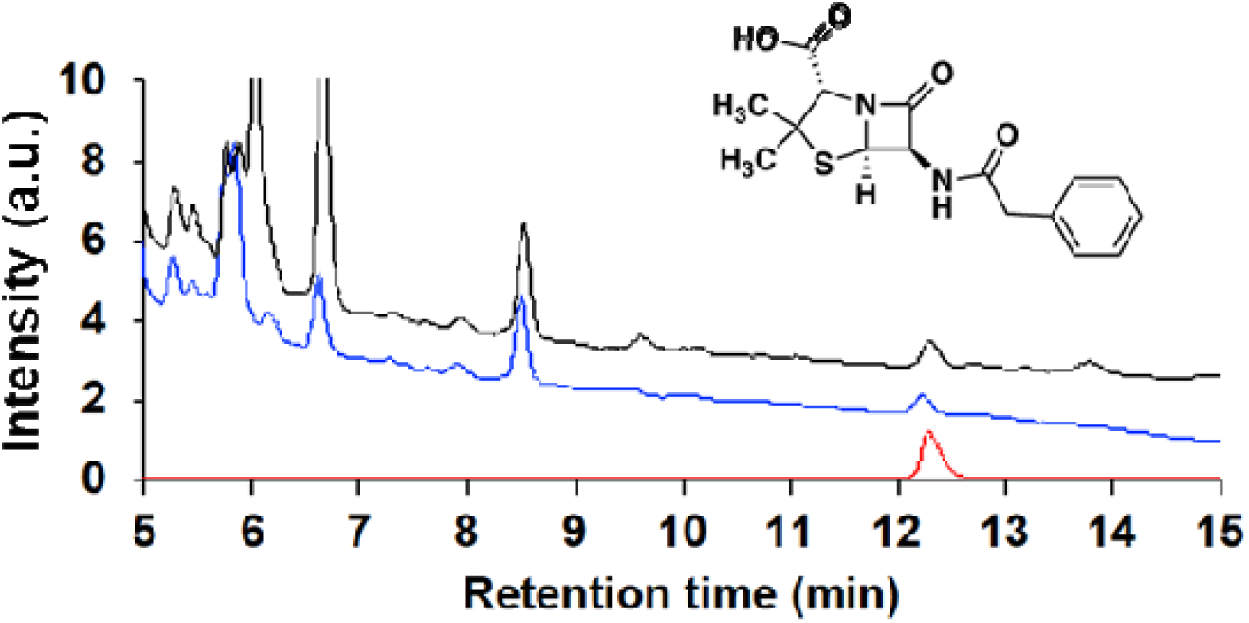
HPLC chromatograms of the reference molecule, commercially obtained penicillin G, and *Penicillium chrysogenum* KF 425 culture extract. Black line: mycelium extract in F7 medium; blue line: supernatant in F7 medium; red line: benzylpenicillin.

### Reference Penicillin G Raman Spectra

A solid powder (Figure 2a) and solution dissolved in PBS (75 mM, Figure 2b) of penicillin G were measured as standard Raman spectra of penicillin G. In these spectra, the Raman bands originating in the β-lactam and thiazolidine rings, the skeleton of the penicillin molecule, and the phenyl ring and specific side chains of penicillin G, are clearly observed. The Raman bands assigned to the β-lactam ring and thiazolidine ring are observed at 850, 950, 1005, 1195, and 1261 cm^−1^;^17^ whereas, those assigned to the phenyl ring are observed at 1005, 1032, 1160, 1585, and 1605 cm^−1^. These bands are observed in both the solid and solution phases, but the solid-phase bands are more clearly observed, having sharper, more well-defined peaks. This is because bandwidth broadening occurs in the solution phase, as a result of collisions and intermolecular interactions.^17^ The Raman band at 1680 cm^−1^ is only observed in the solid-state spectrum. This band is assigned to the C=O stretch of the carboxyl group, which is known to disappear in aqueous solution because of the formation of a carboxylate ion.^18^ In both the solid-phase and the solution-phase spectra, the most obviously apparent penicillin G bands are principally those assigned to the phenyl group. However, in biological samples in general, phenyl groups are widely observed, as components of phenylalanine and tryptophan residues in proteins. Consequently, Raman bands originating from phenyl groups also exist in the Raman spectra of proteins (Figure 2c). Therefore, when the Raman spectra of penicillin G is compared with that of a protein (albumin), there is considerable band overlap. Hence, it is difficult to spectroscopically determine the presence of penicillin G within a mixture containing proteins via the identification of a single intense Raman peak.

**Figure 2.**
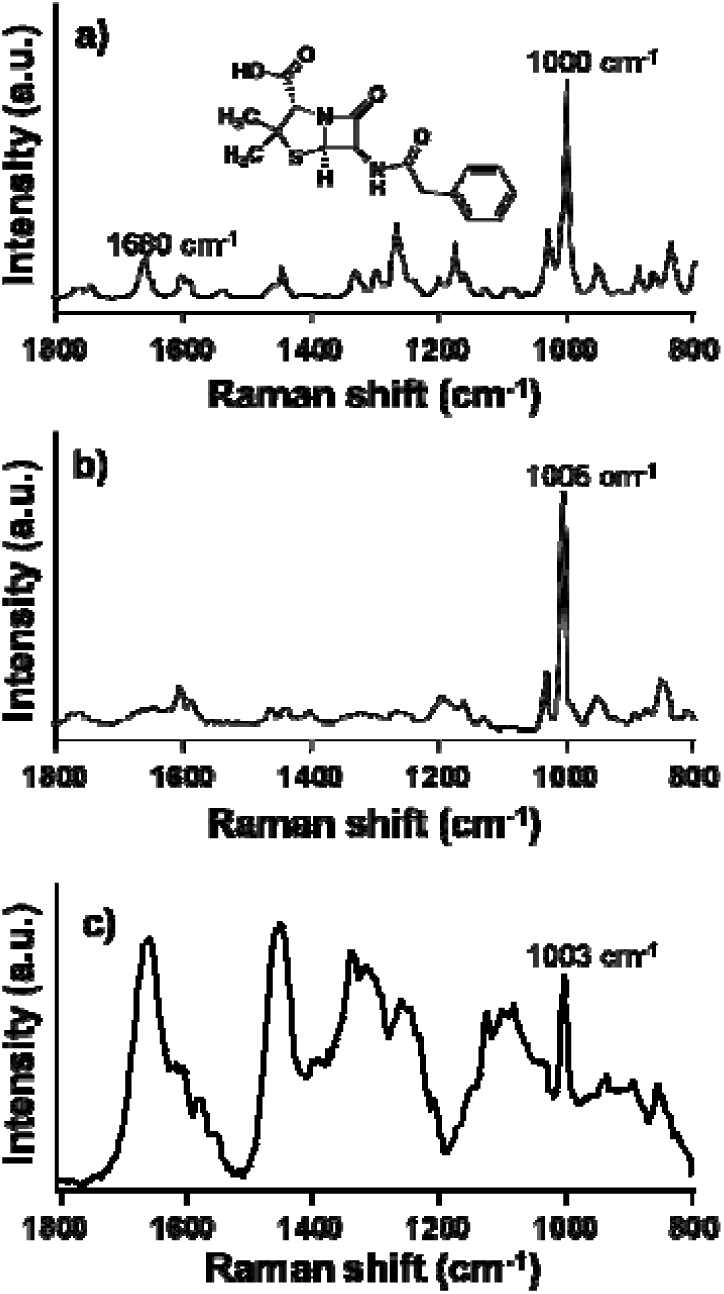
Raman spectra in fingerprint regions and chemical structures of benzylpenicillin potassium: (a) solid benzylpenicillin and (b) benzylpenicillin in PBS solution. (c) For comparison, the Raman spectrum of a protein (albumin) is also given.

### In Situ Detection of Penicillin G in *P. chrysogenum* by MCR-ALS Analysis

In order to accurately evaluate the contribution of penicillin G to the observed superimposed spectra, it is essential to use a multivariate spectral analysis method that evaluates the entirety of each spectrum instead of focusing on a single band. Here, we used a MCR-ALS multivariate analysis technique to verify whether the penicillin G components could be extracted from the measured spectra of *P. chrysogenum* cells. An optimal solution was obtained by MCR-ALS calculation under non-negativity and lasso constraints.^11^ Figure 3 shows the six decomposed spectra produced as the results of MCR-ALS analysis. The Raman spectrum of component 1 is assigned to proteins; its features include peaks at 1004 cm^−1^, the ring breathing mode of phenylalanine and tryptophan residues, 1442 cm^−1^, C–H bending, and 1654 cm^−1^, amide I.^19^ In component 2, Raman bands assigned to α-D-glucan at 942, 1084, 1126, 1208, 1263, 1460 cm^−1^ are apparent.^20^ The polysaccharide constituting the cell wall of filamentous fungi is α-D-glucan, which is a polymer in which D-glucose is linked by glycoside bonds. Component 3 has cytochrome *c* Raman bands at 749, 1128, 1311, and 1583 cm^−1^.^21,22^ Component 4 is assigned to unsaturated lipids, with bands at 1080, 1266, 1301, 1442, 1657, and 1746 cm^−1^.^23^ This can be interpreted as a superposition of the spectra of several cellular lipid constituents. Comparing the band intensity of the C=C stretch at 1657 cm^−1^ with that of the alkyl C–H bend at 1442 cm^−1^, it can be deduced that each lipid molecule contains one or more C=C double bond, on average.^23^ Furthermore, the four peaks observed in the region of 800–1000 cm^−1^ exhibit a pattern consistent with the spectral profile of linoleic acid, which is suggested to be dominant among the various unsaturated fatty acid constituents.^23^ Component 5 has strong Raman bands at 694 cm^−1^ and 1161 cm^−1^, corresponding to polyphosphate P–O–P and PO_2_^−^ stretches, respectively.^24^ Finally, component 6 includes the characteristic Raman bands of penicillin G. The spectrum of component 6 has Raman bands at 1005, 1031, 1160, 1586, and 1602 cm^−1^, which are assigned to the vibrations of the phenyl ring. Furthermore, Raman bands from the β-lactam and thiazolidine rings, and the skeleton of the penicillin molecule, are observed at 849, 1004, and 1193 cm^−1^. The overall spectral pattern of component 6 is identical to the reference Raman spectrum for penicillin G in the solution phase (Figure 2b). In addition, interestingly, a broad Raman band assigned to the O–H bending mode of the water molecule is also observed at 1640 cm^−1^. Hence, in the fungal cell, penicillin G is confirmed to exist in an aqueous-solution state. In summarizing, the MCR-ALS-decomposed spectra shown in Figure 3 demonstrated that we have successfully extracted the Raman spectral components of proteins (1), polysaccharides (2), cytochrome *c* (3), lipids (4), polyphosphates (5), and penicillin G (6). Thus, we have succeeded in separating spectrally overlapped component spectra from the spectrum of a mixture, identifying the spectrum of penicillin G and that of proteins.

**Figure 3.**
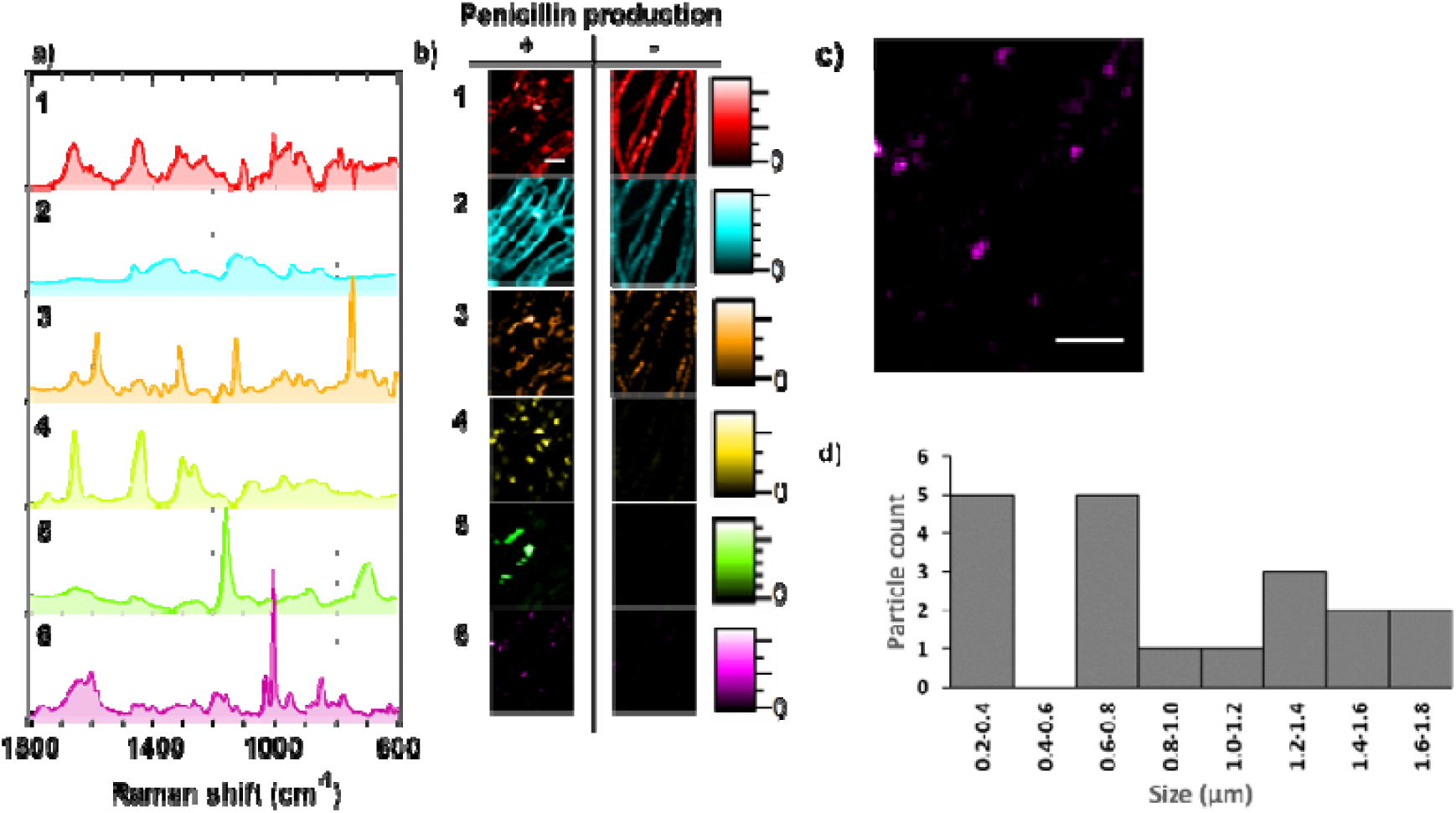
Raman spectra (a) corresponding to (1) proteins, (2) polysaccharides, (3) cytochrome *c*, (4) lipids, (5) polyphosphates, and (6) penicillin G obtained from MCR-ALS analysis and (b) distribution maps for components 1–6 together with overlaid corresponding optical microscopy images and (c) enlarged image of component 6. Scale bar = 5 µm. (d) Particle-size distribution for particles observed in the map of the penicillin component.

From the intensity profile matrix obtained from the MCR-ALS analysis, we reconstructed maps of the molecular distribution within the cells, as shown in Figure 3b. Consequently, we observe that each biological component displays a different localization pattern in the fungal mycelia. From the map for component 1, it is seen that proteins are homogeneously distributed in the cytoplasm of the fungus. In contrast, the component 2 map shows that polysaccharides, as cell-wall components, have higher intensities at the outer edge of the mycelia. The other components, cytochrome *c*, lipids, polyphosphate, and penicillin G, are observed to be localized inside the cells. Cytochrome *c* and lipids are reported to be localized intracellularly and cytochrome *c* is found in mitochondria, and lipids are accumulated in the interior of the fungus, as an energy storage material for its growth. ^25^ In particular, linoleic acid is known to be the most abundant constituent lipid (about 30 %) of the accumulated lipids in *P. chrysogenum*,^26^ in accord with the above-mentioned spectral assignment. *P. chrysogenum* is taken into the body by binding extracellular magnesium and iron ions to sodium polyphosphate, and stored into the body as polyphosphate granules.^27^ Fig 3 (b)-5 shows local accumulation of polyphosphate with various size of granule-like aggregation. The accumulations of lipids and polyphosphate granules seem not to be observed in mycelial cells cultured in TSB medium. However, these components were detected in small amount (Figure S1). F7 medium in which fatty acids and polyphosphates are detected in large amounts contains substrates for metabolites such as sucrose and glycol. On the other hand, these components are not contained in TSB medium. Therefore, only small amounts of fatty acids and polyphosphoric acids are produced in mycelial cells cultured in TSB medium. Finally, the accumulation of penicillin G is only observed in mycelial cells cultured in F7 medium, but not in mycelial cells cultured in TSB medium (Figure 3b).

As described above, we demonstrated that the penicillin G component can be detected by MCR-ALS analysis from the complex and overlapping spectral data sets, and the subcellular localization of penicillin G can also be revealed. In the distribution map for penicillin G, localized particle-like features are apparent. It has been reported that penicillin G is biosynthesized within peroxisomes. It is also known that increases in vacuoles and pexophagy (autophagy of peroxisomes) result in excessive secretion of penicillin.^28^ For the secretion of penicillin G, a mechanism involving exocytosis has been proposed as follows: penicillin G is taken up into vacuoles by pexophagy, and vacuolar vesicles are formed, by vacuolar budding, which fuse to the cell membrane and are finally secreted via exocytosis. According to this hypothesis, penicillin G is contained and localized in compartments (peroxisomes, vacuoles, and vacuolar vesicles) within the mycelial body. The results of our study indicate the localization of penicillin G inside the mycelial cells. The particle sizes in the penicillin G distribution map were estimated by using ImageJ software^29,30^ and can be approximately classified into three size groups: 0.2–0.4 µm, 0.6–0.8 µm, and >1 µm. (Figure 3d). These sizes are consistent with the reported sizes of vacuolar vesicles, peroxisomes, and vacuoles, respectively.^28^ In addition, it was suggested from the MCR-ALS-decomposed spectra that penicillin G is present in the cells as a solute in an aqueous solution, as described above. Hence, from these results, it can be concluded that penicillin G is localized in particles of ∼1-µm in size that have high water contents. This result supports the hypothesis that penicillin G is present in the vacuoles. Thus far, there have been no reports of direct visualization of the localization of penicillin G inside mycelial cells, and the present results therefore provide the most decisive available evidence in support of the exocytosis secretion mechanism. Although the number of measurements in this study is limited and the resolution of the Raman images is not sufficient for more detailed sub-cellular localization observation, further statistical analysis with more measurements using higher-resolution images should provide a clear understanding of the mechanism of penicillin secretion in *P. chrysogenum*.

For a detailed analysis of the penicillin G metabolism in the fungus body, metabolic changes over time were assessed. During *P. chrysogenum* cultivation, cells were collected every 24 h and Raman mapping measurements were conducted. The obtained data were subjected to MCR-ALS analysis, in which the spectral components obtained by the MCR-ALS analysis described above were used as reference spectra. From the MCR-ALS intensity profile matrix, **H**, the contribution and time-course change of each molecular component were evaluated. Here, at each time point, the intensities of each respective component were averaged inside the cells and normalized by the average intensity of the protein component. This is because absolute quantification is difficult, and since proteins are distributed throughout the fungal bodies, the relative change with respect to the protein amount served as the metric for this study. As a result, time-dependent changes in cytochrome *c*, lipids, polyphosphates, and penicillin G were observed (Figure 4). The relative intensity of cytochrome *c* reaches a maximum after 72 h of culture, and tends to decrease until 96 h, and plateaus after 96 h. This intensity change indicates that cytochrome *c* is consumed in the fungus by a metabolic process. The relative intensities of lipids and polyphosphate were negatively correlated. Polyphosphates have been reported to have multiple functions, one of which is that they are known to be a substrate for ATP synthesis. In addition, fatty acids are synthesized by consuming ATP. Therefore, we propose the following hypothesis: In the growth stage, fatty acids are synthesized by ATP generation which is accompanied by polyphosphate degradation. After sufficient fatty acids have been accumulated, polyphosphate decomposition is suspended and the fatty acids are consumed by another metabolism.^32^ Finally, the relative intensity of penicillin G gradually decreases. Therefore, it is suggested that penicillin G is gradually secreted into the medium during the growth of fungi.

**Figure 4.**
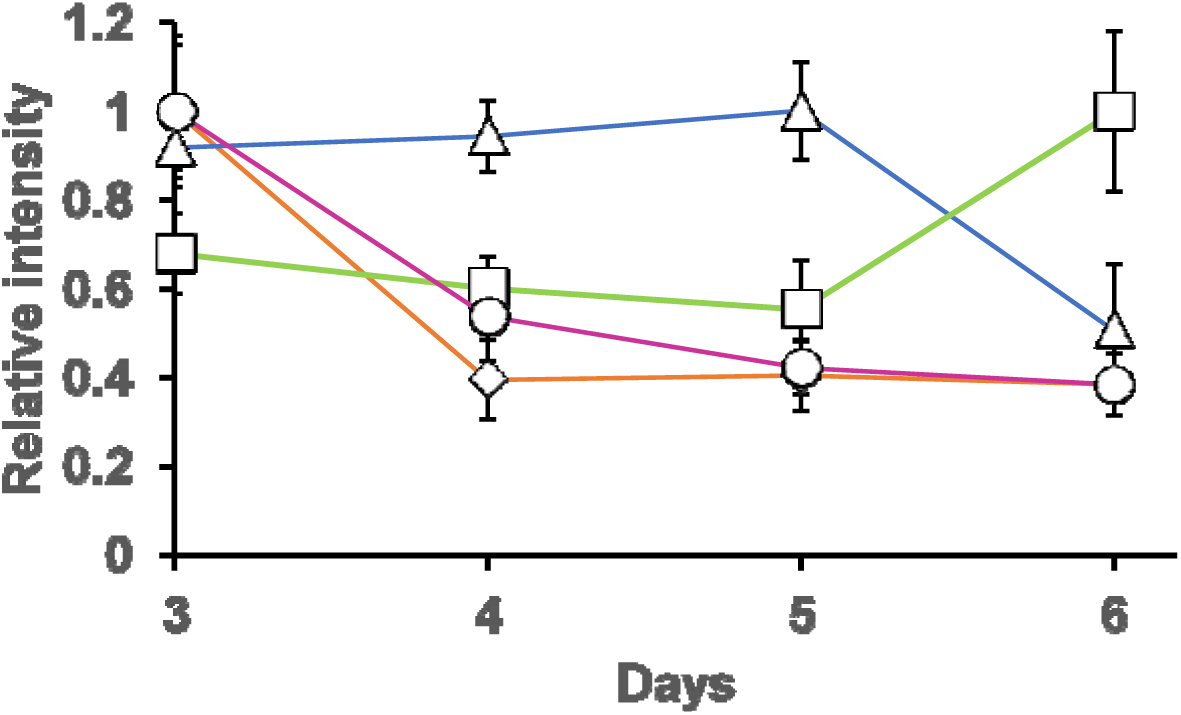
Relative intensity of Raman spectral components corresponding to (⍰) cytochrome *c*, (Δ) lipids, (□) polyphosphates, and (○) penicillin G obtained from MCR-ALS analysis of a hypha cell at each day during cultivation. The intensity of each Raman spectral component is normalized to that of proteins at the same time point.

These results lead us to the conclusion that MCR-ALS analysis can extract biomolecular components from overlapped and complicated spectra and accurately visualize the distributions of biomolecular components. Notably, it is shown that MCR-ALS analysis can disentangle the Raman spectrum of penicillin G and from that of proteins, which has characteristic Raman bands at the same wavenumber positions. By evaluating the spatial distribution of penicillin G inside the fungal bodies, we found evidence that penicillin G is contained in peroxisomes, vacuoles, and vacuolar vesicles, which is evidence in support of the exocytosis-based mechanism for the secretion of penicillin G. Thus, this method has potential for use in analysis of the process of production, accumulation, and secretion of secondary metabolites, together with observation of the molecular dynamics of other cellular constituents. In future, it is expected that this approach will be applied to exploratory research on secondary metabolites, facilitating the acquisition of understanding on the accumulation and secretion process, and in screening for high-producing strains and culture conditions.

## EXPERIMENTAL SECTION

### Fermentation and Sample Preparation

In this study, *P. chrysogenum* KF 425 was used for Raman microspectroscopic analysis. A loop of spores of strain KF 425 was inoculated into 5 mL of the seed medium, consisting of 0.5% polypeptone, 2% glucose, 0.2% yeast extract, and 0.1% KH_2_PO_4_, 0.05% MgSO_4_·7H_2_O. The culture liquid was incubated at 27 °C with shaking at 200 rpm for 2 days to grow the mycelia. The strain KF425 was inoculated into 100 mL of F7 medium, consisting 2% sucrose, 1% glucose, 3% corn steep powder, 0.5% meat extract, 0.05% MgSO_4_·7H_2_O, 0.1% KH_2_PO_4_, and 0.3% CaCO_3_ (adjusted to pH 7.3 before sterilization). The liquid culture was incubated at 27 °C with shaking at 200 rpm for 6 days to produce penicillin G. After 4, 5, and 6 days of cultivation, 1 mL of the culture broth was recovered and separated into mycelium and supernatant by centrifugation. The mycelium was extracted with 1 mL ethanol. TSB medium, containing of 1.7% tryptone, 0.3% Soytone, 0.25% glucose, 0.5% sodium chloride, and 0.25% dipotassium phosphate (adjusted to pH 7.3 before sterilization), was used as negative control for the production of penicillin G under the same culture conditions. Benzylpenicillin potassium (Penicillin G) (Fujifilm Wako Pure Chemical Co., Tokyo, Japan) was used as a reference standard compound.

### Penicillin G Productivity in Antibacterial Activity Test and High-Performance Liquid Chromatography (HPLC)

In order to confirm the penicillin G productivity, antibacterial activity testing and high-performance liquid chromatography (HPLC) were performed. *Kocuria rhizophira* NBRC 103217 was used as a test bacterium for the antimicrobial activity assessment. An equal volume of ethanol was added to the strain KF 425 culture broth, and the supernatant was recovered by centrifugation. The antimicrobial activity was evaluated by a disk-diffusion agar method using paper disks (φ 8 mm) with 10 μL of culture extract. After incubation, the productivity of penicillin G was confirmed by inhibition zone assay. The penicillin G productivity of strain KF 425 was also confirmed by HPLC (Hitachi LaChrom Elite, Hitachi High-Technologies Corp., Japan) using an Ascentis Express C 18 column instrument (25 cm × 4.6 mm, 5 μm, Sigma-Aldrich Corp, St Louis, USA) at 40 °C. For the linear gradient elution, solvent A was ultrapure water with 1% formic acid and solvent B was acetonitrile containing 1% formic acid (A:B, 25:75, v/v). The elution was executed with a flow rate of 1 mL/min for 30 min; the injection volume was 5 μL, and UV detection was carried out using a photodiode array detector. The standard curve was prepared by making serial dilutions, using concentration of 0.2, 0.1, 0.05, 0.025, 0.0125, 0.00625, 0.00313 mg/mL of benzylpenicillin potassium dissolved in PBS (-).

### Raman Microspectroscopy and Imaging

All Raman spectroscopic measurements were carried out by using a laboratory-built confocal Raman microspectroscopy setup, as described in our previous report.^10^ In brief, a 532-nm laser was focused by an objective lens (100×, 1.4-NA) onto a sample placed on the stage of an inverted microscope. The same objective collected the scattered light, and, after appropriate filtering, the Raman spectra were recorded using a spectrometer with a spectral resolution of 3.0 cm^−1^. The spatial resolution of the Raman imaging system was 0.3 × 0.3 μm in the lateral directions and 2.6 μm in the axial direction. For measuring the penicillin G reference spectra, commercially available benzylpenicillin potassium powder (Wako Pure Chemical Industries, Ltd., Osaka, Japan) was dissolved in PBS. The laser power was set to 15 mW, the exposure time to 10 s, and Raman spectral measurements were conducted. The measurements on the *P. chrysogenum* cells were performed as follows. Two milliliters of the culture broth of strain KF 425 were sampled. The fungal mycelia were washed with PBS, placed on a clean coverslip, and sealed with nail polish. Raman spectra of the mycelia were collected over a region spanning 20 × 25⍰μm, with a 0.5-μm scanning interval, and thus 2000 spectra were obtained. Similarly, Raman spectra of single hyphae were collected over a 10 × 15⍰μm region with a 0.33-μm scanning interval, and 1350 spectra were obtained. The laser power was set to 10 mW, and the exposure time was 1 s for each point spectrum. For the time-course measurement, cells were collected every 24 h for Raman mapping measurements.

### Data Analysis

Raman spectral data were preprocessed using IGOR Pro software (WaveMetrics, Inc., Lake Oswego, OR, USA). After wavenumber calibration using the Raman spectrum of indene and intensity correction using the spectrum of a halogen lamp from the microscope, all the spectra acquired in the Raman mapping experiment were combined into a single matrix. Prior to the MCR-ALS calculation, noise reduction by singular value decomposition (SVD) was performed.

Following the data preprocessing, the MCR-ALS calculation was conducted based on the method developed in our previous study. ^11^ MCR-ALS analysis is a method for performing the following optimization calculation iteratively, and it is necessary first to provide initial values.

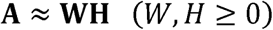

Here, the initial values were set in the matrix **W** from a combination of the SVD output spectra and reference spectra. Thus, from the SVD result, the number of dominant components was determined to be 11, and the corresponding SVD decomposed spectra were used as initial column vectors in the **W** matrix. In addition, the initial **W** matrix was created by concatenating the Raman spectra of the standard antibiotics and background components such as glass and PBS as reference spectra. Among these, only the background spectral components were treated as fixed components, and the ALS iterative calculation provided optimized spectra for the other components. In the ALS optimization process, appropriate physically adaptable constraints make the spectral decomposition results assignable to specific molecular components. Non-negativity is the essential constraint, which is based on the fact that Raman spectra and concentrations of molecules are always non-negative. In this study, the optimal solutions were obtained by introducing an additional constraint, *l*_1_ regularization for the process of estimating **H** from **W**, as follows.

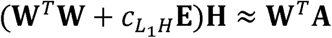

This is known as Lasso regularization, and facilitates the obtaining of sparse solutions.^33^ Physically, this means that each molecule is spatially localized. These physically adaptable constraints are effectively used in MCR-ALS analysis to achieve physico-chemically interpretable spectral decomposition. In this study, the coefficient of the *l*_1_-norm regularization term, 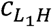, was determined to be 0.001 by cross-validation. Using the above-described MCR optimization, decomposed spectra were obtained from the **W** matrix, and the corresponding distribution maps were obtained by reconstructing the **H** matrix according to the position information from the mapping experiment. These MCR-ALS calculations were performed by a home-built program written in python, using the SciPy library.

## Author Contributions

SH, AM^*§*^, SZ, AT, TN, AM^‖^, YT, and HT conceived and designed the experiments. SH and TN conducted antimicrobial activity test and HPLC method. SH and AM^*§*^ conducted Raman spectroscopic measurements and the data analysis. AM^*§*^ contributed to the analytic tools. SH, AM^*§*^, TN, and HT wrote the manuscript. All authors have given approval to the final version of the manuscript.

## ACKNOWLEDGMENT

This work was supported by a Grant-in-Aid for Scientific Research S (no. 17H06158).

