## Supporting Information for "Detection of Penicillin G Produced by *Penicillium chrysogenum* KF 425 in Vivo with Raman Microspectroscopy and Multivariate Curve Resolution-Alternating Least Squares Methods"

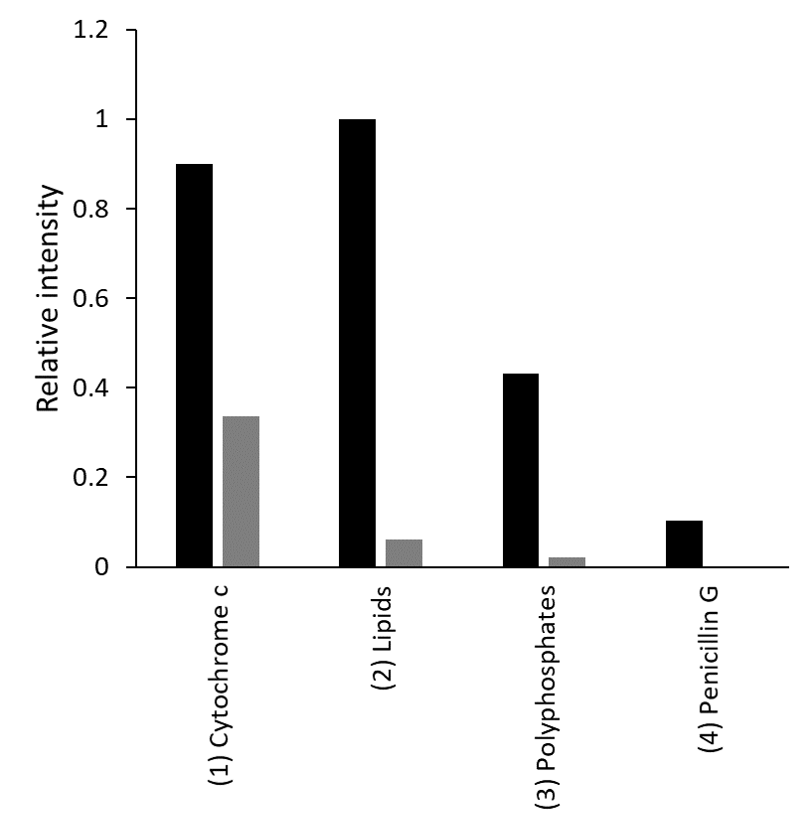

**Figure S1.** Relative intensity (/protein) of Raman spectral components (Figure 3a, b) corresponding to (1) cytochrome c, (2) lipids, (3) polyphosphates, and (4) penicillin G obtained from MCR-ALS analysis of mycelium cells. The intensity of each Raman spectral component is normalized to that of proteins at the same measurement point.
